# Distinct experience-dependent reorganization of rhythmic syllable tracking in 6-month-old term and preterm infants

**DOI:** 10.64898/2026.09.23.753806

**Authors:** Isabelle Rambosson, Damien Benis, Francisca Barcos-Munoz, Claire Kabdebon, Leonardo Ceravolo, Didier Grandjean, Manuela Filippa

**Affiliations:** Neuroscience of Emotion and Affective Dynamics Lab, Department of Psychology, University of Geneva, Geneva, Switzerland; Swiss Center for Affective Sciences, University of Geneva, Geneva, Switzerland; Division of Development and growth, Department of Women, child and adolescent medicine, University Hospital of Geneva, Geneva, Switzerland; Division of Neonatal and Intensive Care, Department of Pediatrics, Gynecology and Obstetrics, University Hospitals of Geneva, Geneva, Switzerland; CRPN, UMR 7077, CNRS, Aix Marseille Université, France

## Abstract

Preterm birth increases the risk of later language difficulties, yet its impact on early neural mechanisms supporting language acquisition, including speech rhythm tracking, remains understudied. Using EEG, we measured neural tracking of delta- and theta-rate syllable sequences produced by the infant’s mother or an unfamiliar speaker in term and preterm 6-month-old infants, modeling how oscillatory responses evolved over the course of a testing session. Term infants progressively strengthened frequency-matched delta tuning over all frontotemporal regions, whereas preterm infants did so only over the right temporal region. Over left frontal channels, this divergence was one of frequency selectivity: term infants became increasingly selective for the delta rhythm, whereas preterm infants showed no reliable selectivity. The frontal group difference emerged only with the unfamiliar speaker, where term infants’ build-up of delta tuning was largely specific. Preterm birth is thus associated with a different refinement of frequency-specific tuning during exposure, rather than with a general weakness in tracking speech rhythm, identifying a candidate neural pathway with implications for early language learning.

## Introduction

Preterm birth, which is defined as delivery before 37 weeks of gestation, affects approximately 10% of births worldwide and constitutes a significant risk factor for neurodevelopmental difficulties^1^. Although survival rates have improved substantially over recent decades, children born prematurely remain at an increased risk for a broad range of adverse outcomes that extend beyond the neonatal period and often persist throughout childhood. Deficits in language development are among the most consistently documented, with preterm children showing delays in vocabulary acquisition, slower phonological processing, and persistent difficulties in receptive language comprehension^2–5^. These language difficulties are frequently associated with social communication and pragmatic difficulties^6–8^, as well as reduced academic achievement, especially in literacy, reading, and mathematics^9,10^. Language and communication vulnerabilities do not emerge in isolation. Indeed, preterm children also exhibit atypical sensory processing^11^, altered multisensory integration^12^, differences in socio-emotional functioning^13^, and difficulties in early parent-infant co-regulation^14,15^.

Although these domains are usually studied independently, each requires integrating information that unfolds over time. Indeed, language processing relies on extracting simultaneous statistical and structural regularities distributed across multiple temporal scales, spanning sub-syllabic properties of phonemes to the rhythmic patterning of syllables and the slower unfolding of prosodic and syntactic structures^16^. Similarly, multisensory processing relies on the temporal alignment of information conveyed through different sensory channels^17^, and perceiving this alignment crucially depends on whether the signals fall within the appropriate temporal integration windows^18^. Furthermore, successful dyadic co-regulation (i.e., the dynamic coordination of affect and behavior between the caregiver and the infant) is a fundamentally rhythmic process that emerges from the mutual adjustment of behavioral, emotional, and communicative signals across time^19^. Tracking, predicting, and integrating the temporal organization of incoming information is therefore a shared functional demand across many of the developmental challenges associated with prematurity.

Speech rhythm provides a particularly powerful and tractable model for investigating temporal processing mechanisms in early development. Natural speech contains inherent hierarchically nested periodicities: syllabic rate falls in the theta range (4–8 Hz), while stress and phrasal patterns operate at slower delta-band rates (1–3 Hz), and these rhythmic levels are cross-coupled in a structured way^16^. Infant-directed speech (IDS), the form of speech naturally addressed to infants, amplifies these cues. It is slower, more melodic, and rhythmically regular than adult-directed speech^20,21^. This is thought to enhance its salience, supporting language acquisition by facilitating attention, segmentation, and prediction ^22–24^. Contemporary neurocomputational models propose that the brain encodes these rhythmic regularities via neural oscillations at different frequencies that entrain to the temporal structure of the speech signal^16,25^. Specifically, delta-band activity supports the parsing of slower prosodic structures^26^, and theta-band activity facilitates syllabic segmentation^27^. Accordingly, neural entrainment, defined as the alignment of endogenous oscillatory activity with external rhythmic input^28^, has emerged as a key mechanism through which infants may parse, predict, and extract linguistic information from the speech stream^29^. Crucially, delta and theta oscillations that support syllabic and phrasal processing have been shown to be present from birth^30^ and to demonstrate developmental differences in cortical tracking across the first year of life^29^, suggesting that the neural architecture for speech entrainment is available from the earliest stages of development and continues to mature with experience.

Early research tended to characterize the preterm brain as immature, with reduced capacity for complex auditory or rhythmic processing. Recent work, however, has substantially revised this picture. Studies using electroencephalography (EEG) have shown that preterm neonates exhibit neural responses that track the rhythmic structure of sound sequences even before reaching term-equivalent age, including evidence of beat tracking and meter encoding^31,32^, as well as temporal prediction abilities by detecting temporal violations in rhythmic sequences and by periodicity encoding at expected beat positions even in the absence of sound^33,34^. These findings show that fundamental capacities for temporal processing are already operative in the preterm brain before term, raising important questions about how these abilities are recruited during speech processing. Despite growing interest in neural entrainment during infancy, relatively little is known about how preterm infants encode the temporal structure of speech and whether this process differs from that of full-term infants.

One factor that may modulate neural entrainment to speech in early infancy is the speaker’s identity, especially whether the voice belongs to the infant’s mother. Fetuses are exposed to the maternal voice throughout gestation, and this exposure has well-documented effects on auditory preferences at birth^35^. In full-term neonates, the maternal voice elicits distinct early frontotemporal neural responses compared with unfamiliar female voices, and this preference is detectable as early as the first postnatal days^36^. For preterm infants, the question is particularly salient: compared to full-term neonates, preterm infants at term-equivalent age show atypical neural processing of voices, recruiting a more distributed frontotemporal network^37^, with a lack of selective neural responses to the maternal voice in the theta frequency band over bilateral temporal regions^38^, suggesting that reduced or disrupted exposure to the maternal voice during a period of active auditory system development may alter the tuning of frontotemporal auditory processing mechanisms, with potential consequences for language acquisition^39^. However, whether voice familiarity modulates neural tracking of rhythmic syllable sequences specifically remains largely unexplored in this population. Because these voice-related responses involve frontal and temporal sites in both term and preterm infants, our analyses center on the left and right frontal (LF, RF) and left and right temporal (LT, RT) regions.

The present study examined neural tracking of rhythmically regular syllable sequences in term and preterm infants at 6 months (corrected age), across repeated presentations within a single testing session (see Figure 1). We refer to this process as neural tracking rather than entrainment, because increased power at the stimulation rate does not by itself establish entrainment (see Limitations). We assessed whether this tracking becomes increasingly frequency-specific in the delta and theta frequency bands as exposure accumulates, and whether speaker familiarity modulates this process. Based on the evidence reviewed above, we derived two predictions. First, building on evidence that prematurity alters the frequency-specific organization of cortical oscillatory networks, particularly in the delta and theta bands^42^, we predicted that term and preterm infants would differ in two complementary indices of oscillatory tuning:

i. *Canonical* tuning (i.e., matched delta stimulation and delta-band response) was expected to differ between groups primarily in the delta band, with stronger delta tuning in term infants: delta-rate modulation is the dominant timescale of infant-directed speech^20,24^ and provides the prosodic markers on which early parsing depends^26^, making it the canonical pair most likely to index mature frequency-specific tracking^43^.
ii. *Cross-frequency* tuning (i.e., mismatched theta stimulation and delta-band response) was expected to be more prominent in preterm infants, reflecting a less differentiated oscillatory organization.

**Figure 1.**
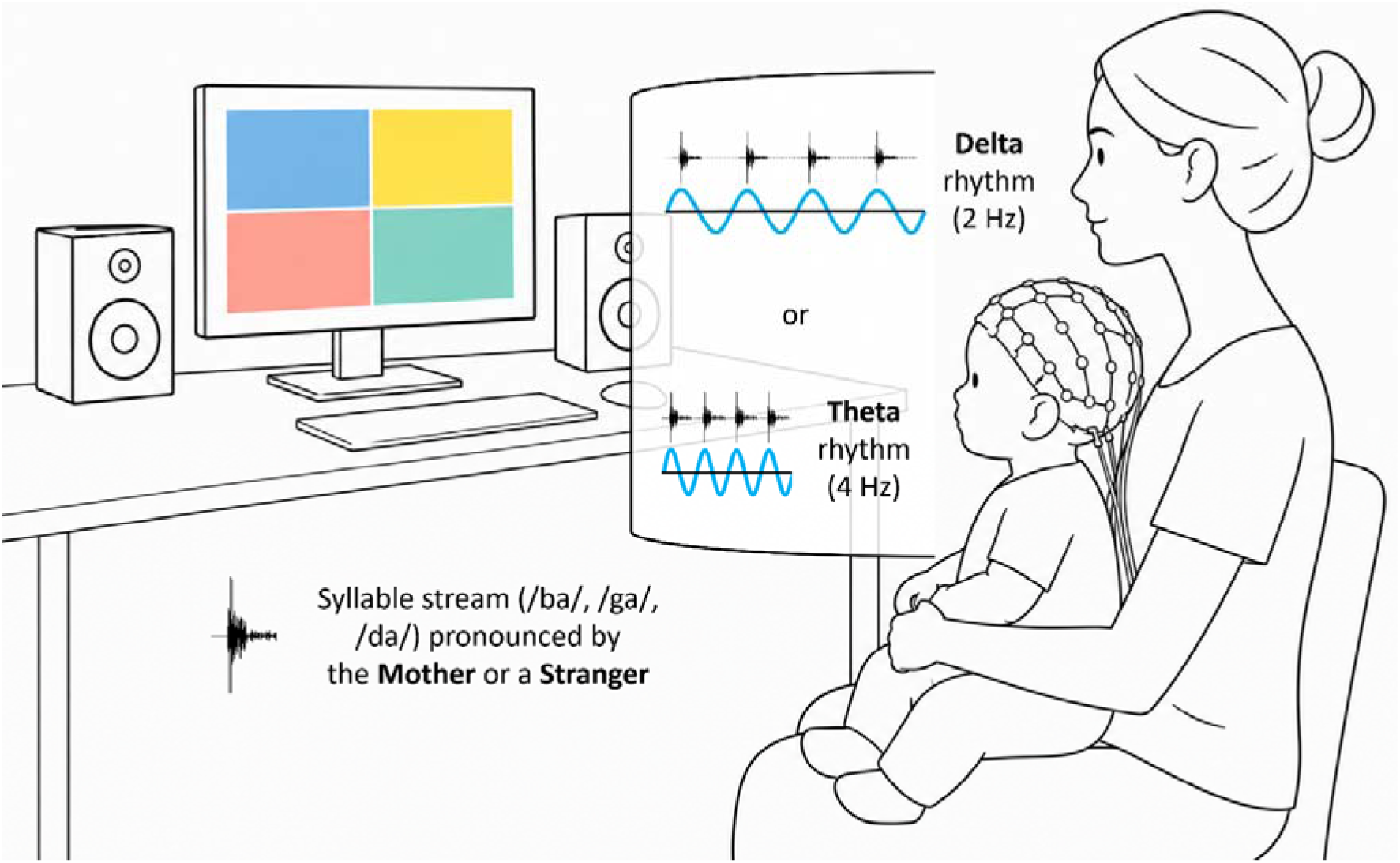
Illustration of the experimental paradigm. Infants listened to four-syllable sequences (/ba/, /ga/, or /da/) spoken by either their own mother or an unfamiliar female speaker, while colored geometric shapes were displayed on the screen. Syllables were delivered every 500 ms (delta rate, 2 Hz) or every 250 ms (theta rate, 4 Hz). The non-rhythmic (NR) control condition is not depicted (see STAR Methods).

Second, we examined whether the maternal (vs. unfamiliar) voice would modulate this term– preterm difference in canonical and cross-tuning conditions. By characterizing how preterm infants track the temporal structure of rhythmic syllable sequences, the study aims to provide new insight into neural mechanisms that may contribute to the developmental trajectories associated with prematurity.

## Results

Rhythm tracking was examined across four frontotemporal regions of interest (ROIs), spanning the left and right frontal and temporal cortices. Within these four, only ROIs showing significant confirmatory effects were carried forward to the exploratory block-wise contrasts. Supporting statistics for each subsection are given in the Supplementary Tables.

### 1. Term and preterm infants did not differ on all background variables other than those defining prematurity

Term (*n* = 30) and preterm (*n* = 17) infants were tested at 6 months of (corrected) age (cf. Table S1). The groups differed only in gestational age and birth weight (both *p* < .0001), the two variables that define prematurity. Corrected age at test, sex distribution, maternal depressive symptoms, parenting stress, infant temperament, and parental socioeconomic status did not differ between groups (all *ps* ≥ .0619; Table S1). Because gestational age and birth weight are confounded with group by definition, they were not entered as covariates.

### 2. Neural tracking changes over the course of exposure in a group-, rhythm-, band-, and region-dependent way

We asked whether neural tracking of rhythmic speech develops differently over the experiment in term and preterm infants.

Power (dB) in the delta (Delta) and theta (Theta) bands was modeled on 10,159,904 observations from 47 infants with a Gaussian generalized linear mixed model (Prediction a; Wald χ² tests, type II). The model crossed stimulation rhythm (delta rate [DEL, 2 Hz], theta rate [THE, 4 Hz], or non-rhythmic [NR]), frequency band, group, block order, and region of interest (ROI), with random intercepts for infant and channel (see STAR Methods). The central (CEN), parietal (LP, RP), and posterior (POS) ROIs entered the same model, and their results are given in Tables S3–S5, S8, S9, S11, S12, and S14.

The change in power across blocks differed between groups depending on rhythm, frequency band, and region (χ²(14) = 51.58, *p* < .0001, and across ROIs χ²(2) = 65.66, *p* < .0001, Table S2), whereas overall power did not differ between groups (χ²(1) = 0.54, *p* = .4624). Group differences therefore lay in how power changed across blocks.

Because the five-way interaction was significant, we decomposed it within each ROI and estimated the linear trajectory of power across blocks (slope, dB per block) for each group, rhythm and band. We then compared term and preterm slopes (Term − Preterm), with *p*-values and simultaneous 95% confidence intervals adjusted for multiplicity across the eight ROIs (mvt adjustment; Figure 2). Slopes within each group are given with unadjusted 95% confidence intervals.

**Figure 2.**
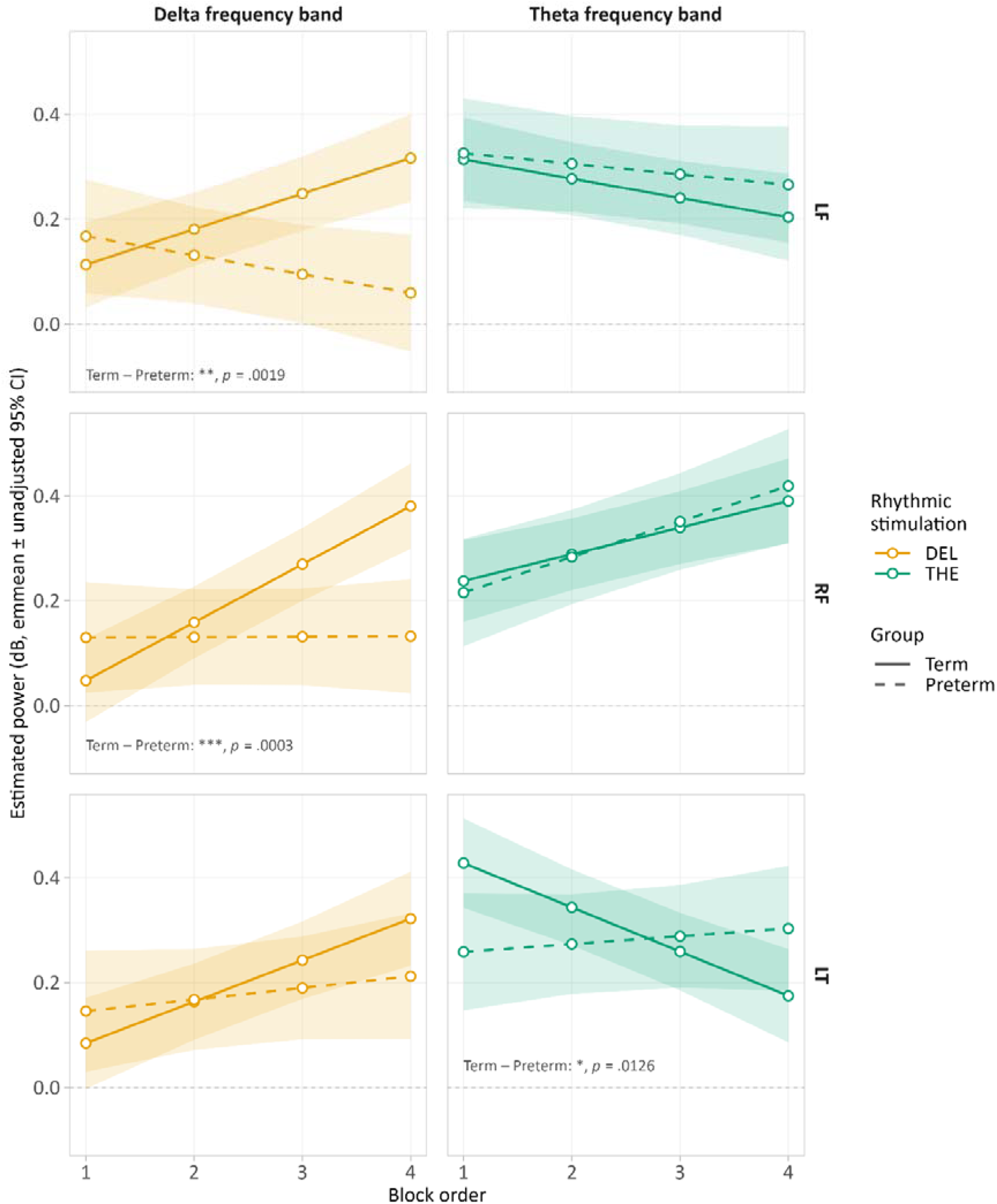
Estimated power trajectories across blocks (1−4) in term and preterm infants for delta and theta tuning in the left frontal, right frontal, and left temporal regions of interest. Each panel shows estimated marginal means (± 95% CI) of the frequency-specific neural tracking (power, in dB) across the four blocks of the experimental session, for the delta stimulation/Delta (orange, left column) and theta stimulation/Theta (green, right column) tuning conditions, and for term (solid line) and preterm (dashed line) infants, in the left frontal (LF), right frontal (RF), and left temporal (LT) regions of interest (rows). Annotations indicate the significance level and p-value of the Term − Preterm difference in slope across blocks when significant (mvt-adjusted across the eight regions of interest). * *p* < .05, ** *p* < .01, *** *p* < .001.

### 3. Term infants increase delta-band power under delta-rate stimulation, with a different topography in preterm infants

Under delta-rate stimulation, the delta-band response (power in the delta band) rose across blocks in term infants in all four frontotemporal ROIs (LF: 0.068, RF: 0.111, LT: 0.079, RT: 0.062 dB per block; Figure 2). Preterm infants showed a different profile: their delta-band response increased only in RT (0.119 dB per block, 95% CI [0.072, 0.166]) and did not change in LF, RF, or LT.

Under the same stimulation, the increase in the delta-band response across blocks differed reliably between term and preterm infants in the two frontal ROIs (Term − Preterm slope difference, multiplicity-adjusted): LF (0.104 [0.023, 0.185], *p* = .0019) and RF (0.110 [0.033, 0.187], *p* = .0003). The temporal ROIs did not differ (LT: 0.057, p = .7529, RT: −0.057, *p* = .7646, Table S3).

The group difference in the delta-band response was not present for non-rhythmic stimulation (NR): delta-band slopes did not differ between term and preterm infants in any ROI (all *ps* ≥ .0560, Table S3). For theta-rate stimulation, the delta-band response differed between groups in only two ROIs, with opposite signs: LF (Term − Preterm: −0.089, *p* = .0103) and RF (0.097, *p* = .0011). In LF, this difference reflected a decrease in term infants (−0.045) and an increase in preterm infants (0.044), whereas in RF it reflected a decrease in preterm infants (−0.091) with no change in term infants (0.007). Thus, in the frontal ROIs, delta-rate stimulation was associated with a larger increase of the delta-band response in term than in preterm infants in both LF (0.104) and RF (0.110), whereas theta-rate stimulation produced a difference of the opposite direction in LF (−0.089) and of the same direction in RF (0.097).

### 4. Theta-band group differences are not specific to theta-rate stimulation

We next asked whether term and preterm infants also differed in theta-band tracking of theta-rate stimulation. In term infants, theta power under theta stimulation decreased across blocks in LF (−0.037 dB per block) and LT (−0.084) and increased in RF (0.051) and RT (0.053). Preterm infants showed increases in RF (0.068) and RT (0.061) and no change in LF or LT.

Group slopes under theta stimulation differed in LT (Term − Preterm: −0.099 [−0.187, −0.011], *p* = .0126) but not in LF, RF, or RT. The same LT difference appeared under delta-rate stimulation (theta-band slope under DEL: −0.109, *p* = .0063), where preterm infants increased (0.109), and term infants did not change (0.000). Theta-band group differences were therefore not tied to theta-rate stimulation, showing a lack of specificity in the tracking response.

### 5. Delta-band selectivity for the matching rhythms is present in all frontotemporal ROIs in term infants but only in RF and RT in preterm infants

The slope differences above do not show whether a band tracks its own rhythm more than the other one. To test this, we computed a selectivity index within each band: the slope under the matching rhythm minus the slope under the mismatching rhythm (Delta band: slope under delta-rate stimulation minus slope under theta-rate stimulation; theta band: slope under theta-rate stimulation minus slope under delta-rate stimulation). Positive values indicate rhythm-specific tuning. We then compared the index between groups (Figure 3 and Table S5).

**Figure 3.**
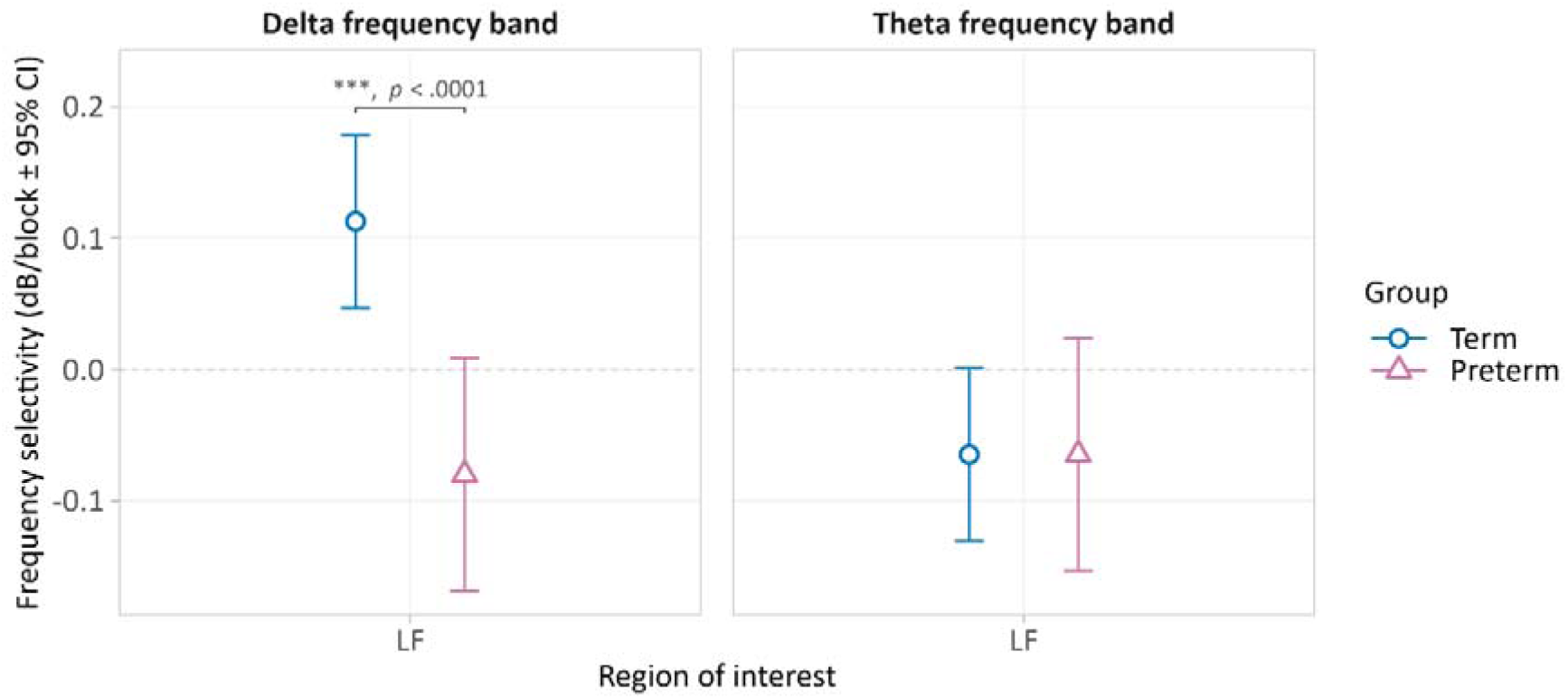
Frequency selectivity in term and preterm infants, by frequency band, in the left frontal region of interest. The selectivity index was computed as the difference between the matched and the mismatched block-order slope within a given response band (Delta band: delta-rate minus theta-rate stimulation, Theta band: theta-rate minus delta-rate stimulation). Since the index contrasts two slopes, it characterizes the change in tuning over the course of the session. Positive values indicate that the difference between the matched and the mismatched response widened across blocks, that is, that tuning became increasingly selective, whereas negative values indicate the converse. Points show the estimated selectivity index for term (blue circles) and preterm (pink triangles) infants in the left frontal region of interest (LF), the only frontotemporal region in which the index differed between groups, together with 95% simultaneous confidence intervals. The bracket marks a significant difference in the selectivity index between the two groups (mvt-adjusted across the eight regions of interest). *** *p* < .001. Full statistics are reported in Table S5.

In the delta band, term infants showed positive selectivity in all four ROIs (LF: 0.113 [0.047, 0.178], RF: 0.104 [0.042, 0.166], RT: 0.112 [0.038, 0.187]; all *ps* ≤ .0001; LT: 0.092 [0.017, 0.166], *p* = .0043). Preterm infants showed it in RF (0.092 [0.007, 0.176], *p* = .0214) and RT (0.173 [0.072, 0.274], *p* < .0001), but not in LF (−0.080, *p* = .1220) or LT (−0.006).

The groups differed in delta-band selectivity in LF (0.193 [0.082, 0.303], *p* < .0001). They did not differ in LT (0.098, *p* = .3009), RF (0.013), or RT (−0.061). RF selectivity was thus comparable across groups despite differences in raw DEL slope.

In the theta band, selectivity was not reliably positive in any frontotemporal ROI. In term infants, it was negative in LT (−0.085, *p* = .0127), and no frontotemporal ROI showed selectivity in preterm infants. No ROI showed a group difference (all *ps* ≥ .7465). The theta band therefore showed no rhythm-specific tuning in either group.

### 6. Speaker familiarity modulates delta-band trajectories differently in term and preterm infants

We next asked whether the maternal voice (Mother) and an unfamiliar voice (Stranger) shaped these trajectories. Adding VOICE improved fit (AIC lower by 1,271 than Prediction a), and voice had a main effect (χ²(1) = 90.27, *p* < .0001). The change in power across blocks differed between groups depending on voice, rhythm, frequency band, and region (χ²(14) = 56.55, *p* < .0001, Table S2).

For delta-rate stimulation, the frontal group difference depended on the voice. With the Stranger’s voice, term infants showed stronger delta increases than preterm infants in LF (0.131 [0.008, 0.254], *p* = .0243) and RF (0.154 [0.036, 0.271], *p* = .0009; Figure 4 and Table S4). These effects arose from preterm slopes close to zero in LF (−0.011) and RF (−0.012), against positive term slopes (LF: 0.120, RF: 0.142). The temporal ROIs did not differ for the Stranger’s voice (LT: 0.045, RT: −0.005).

**Figure 4.**
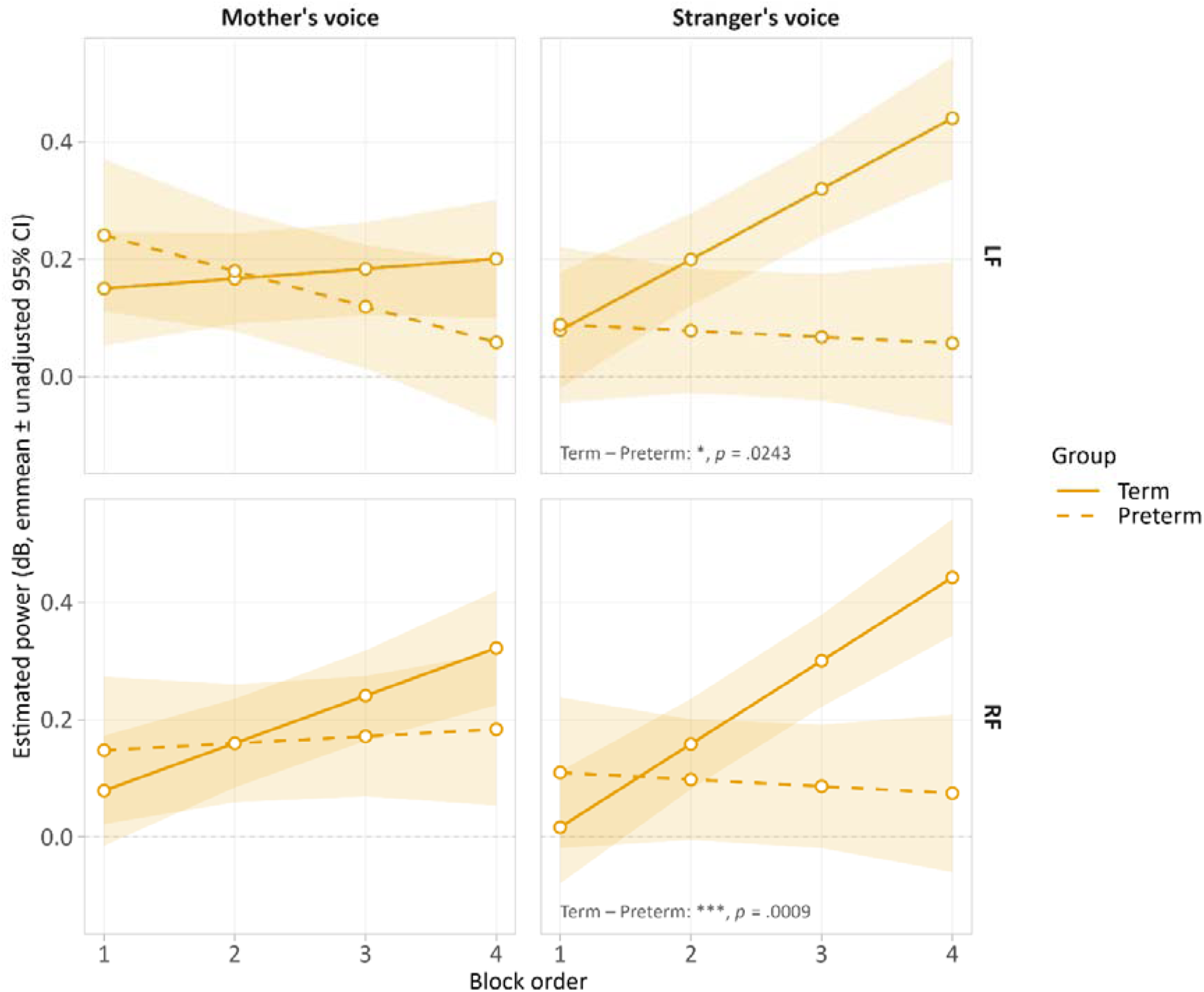
Estimated power trajectories across blocks (1−4) in term and preterm infants for delta tuning in the left frontal (LF) and right frontal (RF) regions of interest, separately for the mother’s voice and stranger’s voice conditions. Each panel shows estimated marginal means (± 95% CI) of the frequency-specific neural tracking (power, in dB) across the four blocks of the experimental session in the delta stimulation/Delta condition, for the mother’s voice (left column) and stranger’s voice (right column) conditions, and for term (solid line) and preterm (dashed line) infants, in the left frontal (LF) and right frontal (RF) regions of interest (rows). Annotations give the significance level and p-value of the Term − Preterm difference in slope across blocks when it is significant (mvt-adjusted across the eight regions of interest). * *p* < .05, *** *p* < .001.

With the mother’s voice, no frontotemporal group difference was significant (LF: 0.078, RF: 0.069, LT: 0.068, RT: −0.106; all n.s.).

To focus on the delta-band response to delta-rate stimulation, we repeated the analysis on these data alone. The group difference in the effect of voice on the change across blocks was again significant (χ²(7) = 42.93, *p* < .0001 across ROIs; χ²(1) = 21.54, *p* < .0001, Table S10), and the frontal results were confirmed: with the Stranger’s voice, term infants showed a larger increase than preterm infants in LF (0.133, *p* = .0075) and RF (0.158, *p* = .0002, Table S11). Block by block, the group difference in the effect of voice was significant in LT (block 2: −0.255, *p* = .0061; block 3: −0.236, *p* = .0296) and in RT (block 1: 0.348, *p* = .0303; block 2: 0.242, *p* = .0126), where it shrank to 0.03 by block 4 (Table S12 and Figure S1). Because these estimates derive from linear trends across blocks, we interpret the overall trend rather than individual blocks.

### 7. The group differences are not explained by other covariates

To exclude confounds, we tested twelve candidate covariates by likelihood-ratio tests against the base models. Block order was the only one that improved fit (Prediction a: χ²(1) = 19.07, *p* < .0001, Prediction b: χ²(1) = 16.89, *p* < .0001). None of the other eleven candidates did (all *ps* = .2271–.9373 across both predictions). Gestational age and birth weight, the only background variables that differed between groups, are confounded with group by definition and were therefore not used as covariates.

Overall, term infants increased delta-band power under delta-rate stimulation in all frontotemporal ROIs and showed matching-rhythm selectivity in each. Preterm infants showed this selectivity only in RF and RT, and increased their delta-band power responses only in RT. They differed from term infants in the frontal ROIs (LF, RF), a difference observed with the unfamiliar voice. Cross-frequency tuning was more prominent in preterm than in term infants in LF and LT, as predicted, but not in RF. Theta-band effects showed no selective tuning in either group.

## Discussion

Preterm birth increases the risk of later language difficulties, yet little is known about how it affects the early neural mechanisms that support language acquisition, such as tracking the rhythm of speech. To better understand this, we investigated how the infant brain tunes to rhythmic speech with exposure, and whether this process differs between term and preterm infants. Our main findings are as follows.

First, overall tracking strength did not differ between groups; the groups differed in how tracking changed across the session. Term infants progressively built up canonical delta tuning over bilateral frontotemporal regions, whereas preterm infants did so only over the right temporal region. Second, the form of the group difference varied by region: over left frontal channels, term infants became increasingly selective for the delta rhythm, whereas preterm infants showed no reliable selectivity. Over right frontal and left temporal channels, the groups did not differ in selectivity but in how a band’s response changed irrespective of the rhythm played: preterm infants showed a less positive delta-band trajectory over right frontal channels and a positive theta-band trajectory over left temporal channels. Third, the frontal difference emerged only with an unfamiliar speaker. Together, these results indicate that preterm birth is associated with a different refinement of frequency-specific tuning during exposure, a candidate neural pathway for early language learning.

### Preterm birth alters how neural tracking is refined with exposure, not its overall level

As predicted, term and preterm infants differed in how neural tracking developed across the session, not in how strongly they tracked speech rhythm overall. Under delta-rate stimulation, term infants increased their delta-band response in all four frontotemporal regions, whereas preterm infants did so only over the right temporal region. No group difference emerged under non-rhythmic stimulation, so the effect reflects tuning to the rhythm and not a general drift in responsiveness. The preterm profile is therefore more circumscribed and less frontal rather than absent.

That a neural response can be reshaped by repeated exposure over the course of a single session, and not only over weeks or months, is already documented in the infant brain. Cortical tracking of the structure embedded in a syllable stream — a stimulus close to the one used here — emerges within roughly two minutes of familiarization even in sleeping neonates, without attention and without instruction^44^. By the age we tested, even which frequency band carries the response can be shifted by recent input, and shifted quickly: brief rhythmic priming with structured sequences selectively enhances subsequent delta- or theta-band tracking of speech in 6- and 10-month-olds, depending on the structure of the priming input^45^. Our design does not identify the process behind these within-session gains. Growing familiarity with the syllables, response enhancement through repetition, fluctuations in alertness, and increasingly precise expectations about syllable timing would all predict them, and we cannot exclude that attention or arousal evolved differently in the two groups.

### Group differences take different forms across regions

The clearest expression of the group differences was a change in frequency selectivity. Over left frontal channels, term infants became increasingly selective for the delta rhythm: their delta-band response to delta-rate stimulation increased while decreasing to theta-rate stimulation relative to preterm infants — two significant effects of opposite sign in the same response band. Preterm infants showed no reliable selectivity: their delta-band response did not rise under delta-rate stimulation and instead rose under theta-rate stimulation, the rhythm that did not carry it. This is not a difference in how much the two groups responded, but in how specific the response was, consistent with less differentiated oscillatory organization our second prediction anticipated. Such sharpening is the signature that experience-dependent tuning is expected to leave: interactive auditory experience sharpens cortical oscillatory responses in infancy^46^, and these gains generalize to speech^47^. We observed such a shift toward the stimulating rhythm across left frontal channels within a single session in term infants, but not in preterm infants. Cross frequency tuning itself was more prominent in preterm infants over left frontal and left temporal channels, but not over right frontal channels, so that our second prediction was only partly supported. That the effect was frontal rather than temporal accords with the distributed frontotemporal network that voice processing recruits in this population^37^.

Elsewhere, the two groups diverged without any group difference in selectivity. Over right frontal channels, the two conditions shifted in parallel: both groups became more frequency-selective across the session to a comparable degree, with term infants increasing their Delta-band response relative to preterm under both delta-rate and theta-rate stimulations, which reflected an increase under delta-rate stimulation in term infants and a decrease under theta-rate stimulation in preterm infants. The selectivity index confirms that the groups did not differ in this region (see Table S5). This is broadly compatible with proposals that early language acquisition depends on the coordination of delta- and theta-scale processing rather than on either alone^48,49^ and with evidence that prematurity alters the frequency-specific organization of cortical oscillatory networks^42^. Theta-band canonical tuning, for which no directional prediction was made, showed no group difference over either frontal region.

These null results should be read in light of the rates used. In 4- and 8-month-olds, tracking of syllable sequences is robust at the syllable rate but inconsistent at a slower rate corresponding to bisyllabic feet^50^; here, both rates were syllabic, which may have favored the slower one. Over left temporal channels, the term theta-band response declined across the session while preterm infants showed no significant trend — a difference in trajectory that is not, as shown below, one of selectivity. The term infants’ decline was not general habituation: their theta-band response declined under theta-rate stimulation while remaining stable under delta-rate stimulation, a rhythm-specific change that a uniform loss of responsiveness could not produce. It also connects the left temporal pattern to the left frontal one. Over left frontal channels, preterm infants extended their delta-band response to the rhythm that did not carry it, a genuine group difference in selectivity. Over left temporal channels, no such difference was present: the theta-band selectivity index was negative in term infants and did not differ between groups (cf. Table S5). What distinguishes the groups there is the overall weighting of the theta-band response, not its tuning. This preserved theta-band selectivity contrasts with a different report in preterm infants, who at term-equivalent age lack the selective theta-band response to the maternal voice that term infants show over bilateral temporal cortex^38^; whether the two findings reflect the same underlying process, given the different contrasts and ages involved, is not settled by our data. Over right temporal channels, no group difference in tuning trajectories reached significance, although preterm infants showed an increasing and selective delta-band response in this region.

The absence of frontal tuning change over frontal channels may relate to the atypical auditory environment of neonatal intensive care. Preterm infants receive substantially less language exposure than they would have received in utero^51^, and the intensive care environment may tune them toward the ambient noise and unpredictable sounds that dominate it at the expense of linguistic input^39^, even though infant-directed speech is increasingly incorporated into developmental care practices. Consistent with this, preterm infants already process the maternal voice atypically at term-equivalent age^38^, and altered neural tuning to voices — but not to faces — is still detectable in preterm preschoolers ^52^. The functional stakes of this circuitry are established independently: in children, the strength of maternal voice-selective responses predicts social communication ability ^53^. Critically, this pathway is modifiable: increasing maternal speech exposure in the neonatal intensive care unit alters white matter connectivity in infants born preterm^54^.

### The group difference emerges with an unfamiliar voice

Over both frontal regions, the canonical delta-tuning difference between groups emerged with an unfamiliar speaker and widened across the session, whereas it was no longer significant with the mother’s voice. This was not because preterm infants caught up with the mother’s voice: over left frontal channels, term infants did not increase their delta tuning either, and over right frontal channels their increase was smaller than with the unfamiliar speaker. In term infants, then, the frontal build-up of delta tuning was largely specific to the unfamiliar voice.

We interpret this in terms of what each speaker demands of the listener. The maternal voice is available to the infant as an already-consolidated representation, shaped by exposure that begins in utero^35^ and elicits distinct frontotemporal responses within the first postnatal days^36^; tracking it requires little new learning. An unfamiliar speaker offers no such representation and must be modeled online, over precisely the timescale on which our blocks unfold.

The familiar voice thus masks, and the unfamiliar voice reveals, a difference in how preterm infants tune to speech rhythm, suggesting that what is altered after preterm birth is the capacity to refine tuning to new input, not tracking itself.

The present design does not test this directly. It does, however, extend earlier work showing atypical responses to the familiar voice in preterm infants at term-equivalent age^37,38^, by 6 months the group difference has shifted to the unfamiliar voice, where the demand is not to recognize a speaker but to build a representation of one. At this age, the unfamiliar voice could reveal a difficulty that familiar input conceals. The early postnatal auditory environment may provide a complementary account. During hospitalization, preterm infants are exposed to numerous unfamiliar voices, including speech exchanged among healthcare staff that is not directed at the infant, whereas term infants cared for at home receive a greater proportion of speech from familiar caregivers. Such exposure, frequent but seldom contingent, may be insufficient to support the refinement of tuning to novel voices and could thereby contribute to the group difference observed here.

### Potential implications for early language acquisition

These findings bear on how speech rhythm scaffolds early language acquisition. Speech is processed across a hierarchy of oscillatory timescales, with delta-band activity tracking prosodic and phrasal structure and theta-band activity tracking syllabic structure, and acquisition is thought to depend on their coordination rather than on either band alone^48,49^. Infant-directed speech is adapted to this hierarchy, carrying stronger delta-range modulation and enhanced delta–theta phase synchronization than adult-directed speech^29,56^, and delta-band entrainment supplies the temporal landmarks on which segmentation and prosodic parsing depend^29,57–59^. Delta tuning that does not consolidate therefore concerns the dimension the infant’s everyday input supplies most redundantly. This is not a loss of rhythmic competence in general: preterm neonates detect rhythm violations, encode beat and meter before term-equivalent age, and show periodicity encoding at expected beat positions even in the absence of sound^32–34^. What develops later across the last trimester is the tracking of hierarchically nested structure rather than the basic beat^31^ — the level built through experience is the level our data show to be compromised. This may bear on the language phenotype of children born preterm, in whom receptive difficulties are among the most consistently documented outcomes^3,4,60^. Receptive language depends on extracting prosodic regularities across extended temporal windows, and cortical tracking dynamics in the first year already predict vocabulary at two years^61^. That link is not merely correlational: in a randomized trial delivering contingent caregiver infant-directed voice to infants born preterm, gains in neural speech-sound differentiation mediated better language outcomes at two years^55^. We did not follow our infants longitudinally, so whether the tuning differences we observed predict later language remains to be tested.

### Limitations

Some limitations qualify the interpretation of these findings. First, the preterm group was small and skewed towards the milder end of the spectrum (*n* = 17) and predominantly late preterm (13 were late preterm, three moderately preterm, and one very preterm). This group composition, together with the exclusion of infants with significant brain injury, suspected neurodevelopmental disorder, or adverse maternal history, yielded a clinically homogeneous, low-risk group that did not differ from term infants on any background variables other than gestational age and birth weight. The group differences reported in this study are therefore unlikely to reflect differences in background characteristics between cohorts. The counterpart of this homogeneity is that findings pertain to low-risk prematurity and cannot be generalized to more severe forms. Second, although the overall sample is comparable to those of infant EEG studies in this field, the small preterm group limits the precision of between-group estimates, particularly for higher-order interactions, whose effects were small. These effects should therefore be treated with caution until they are replicated in a larger cohort. Third, the design is cross-sectional. A single session at 6 months of corrected age allows no inference about individual developmental trajectories or the direction of group differences.

Longitudinal data are needed to establish whether these patterns persist, resolve, or widen with age. Fourth, the present measures rest on spectral power at the stimulation rates, which is why we refer to neural tracking rather than entrainment. This pattern is consistent with frequency tagging but does not demonstrate oscillatory entrainment in the strict sense, which would also require phase alignment with the stimulus and oscillatory activity persisting after its offset^40,41^. Designs including silent post-stimulus periods and phase-based measures would be needed to establish it. Fifth, delta-and theta-rate sequences differed in presentation rate and analysis-window length, and the second harmonic of the 2 Hz rate falls within the theta band. Part of the cross-frequency effects, particularly the theta-band response to delta-rate stimulation, may therefore reflect a harmonic response or a general difference between slower and faster stimulation rather than band-specific tracking. Including additional stimulation rates in future designs would allow these accounts to be separated.

## Conclusions

These findings identify a candidate neural mechanism linking preterm birth to later language and social communication difficulties: not a general weakness in processing speech rhythm, but a failure to refine that processing as exposure accumulates. The term infant brain actively retunes itself to the rhythms it encounters, moment by moment within a single listening session; after preterm birth, this retuning falters, specifically when the voice is unfamiliar — precisely the situation infants must adapt to as their world extends beyond direct experience. Because this kind of experience-dependent tuning is thought to scaffold the syllable- and word-level representations that language later depends on, a disrupted trajectory at 6 months offers a measurable, mechanistic bridge between an early risk due to prematurity and its long-term clinical consequences. More broadly, these results suggest that what may matter for early language learning is not simply whether the infant brain tracks speech rhythm, but whether that tracking can be refined over the course of exposure — a capacity that, in this sample, appeared to be affected by prematurity.

## STAR METHODS

### Key Resources Table

| REAGENT or RESOURCE | SOURCE | IDENTIFIER |
|---|---|---|
| **Deposited data** |  |  |
| Python preprocessing pipeline and R analysis scripts | This paper | https://doi.org/10.26037/yareta:4dtcmgkw4jdwpmawpzcdt4byrq |
| **Software and algorithms** |  |  |
| Python v3.12.3 | Python Software Foundation | https://www.python.org |
| MNE-Python v1.9.0 | Gramfort et al.<sup>62</sup> | https://mne.tools |
| Python-picard v0.8 | Ablin et al.<sup>63</sup> | https://pierreablin.github.io/picard |
| Audacity v3.3–3.7 | Audacity Team<sup>64</sup> | https://www.audacityteam.org |
| Librosa v0.10.2 | McFee et al.<sup>65</sup> | https://librosa.org |
| Praat Vocal Toolkit | Corretge<sup>66</sup> | https://www.praatvocaltoolkit.com |
| Praat v6.3.10 | Boersma and Weenink<sup>67</sup> | https://www.praat.org |
| PsychoPy v2023.2.3 | Peirce et al.<sup>68</sup> | https://www.psychopy.org |
| R v4.5.1 | R Core Team<sup>69</sup> | https://www.r-project.org |
| GlmmTMB v1.1.13 | Brooks et al.<sup>70</sup> | https://CRAN.R-project.org/package=glmmTMB |
| Car v3.1.3 | Fox and Weisberg<sup>71</sup> | https://CRAN.R-project.org/package=car |
| Emmeans v2.0.0 | Lenth and Piaskowski<sup>72</sup> | https://CRAN.R-project.org/package=emmeans |
| Performance v0.15.3 | Lüdecke et al.<sup>73</sup> | https://CRAN.R-project.org/package=performance |
| DHARMa v0.4.7 | Hartig<sup>74</sup> | https://CRAN.R-project.org/package=DHARMa |
| Ggplot2 v4.0.0 | Wickham<sup>75</sup> | https://CRAN.R-project.org/package=ggplot2 |
| **Other** |  |  |
| HydroCel Geodesic Sensor Net, 128 channels | Electrical Geodesics, Inc. | GSN-HydroCel-128 |
| eego amplifier | ANT Neuro | N/A |
| EGI amplifier | Electrical Geodesics, Inc. | N/A |
| Edinburgh Postnatal Depression Scale | Murray and Carothers<sup>76</sup> | N/A |
| Infant Behavior Questionnaire, revised | Gartstein and Rothbart<sup>77</sup> | N/A |
| Parenting Stress Index | Abidin<sup>78</sup> | N/A |
| Socioeconomic status scale | Largo et al.<sup>79</sup> | N/A |

### Experimental Model and Study Participant Details

#### 1.1. Participants

A total of 52 6-month-old infants took part in this study: 33 born at term and 19 born preterm. For preterm infants, age was corrected for prematurity so that all infants were tested at the same developmental age. Detailed demographic and clinical characteristics of both groups are reported in Table 1. The following exclusion criteria were applied: (i) the presence of a clinically significant brain injury, defined as periventricular leukomalacia (grade ≥ 2) or intraventricular hemorrhage (grade ≥ 3); (ii) clinical suspicion of a neurodevelopmental disorder, such as autism spectrum disorder; or (iii) a maternal history of substance abuse during pregnancy or of major depressive episodes.

**Table 1.** Demographic and clinical characteristics of term and preterm experimental groups. ^a^ Missing value imputed via linear regression on gestational age within group. ^b^ Missing values imputed using the group median.

| Participants | Full-term<br>*n* = 30 | Preterm<br>*n* = 17 |
|---|---|---|
| Gestational age (GA) at birth (weeks), *M* ± *SD* | 39.53 ± 1.31 | 34.49 ± 1.90<br>Very preterm (*n* = 1): 29.00 (SD not applicable)<br>Moderate preterm (*n* = 3) : 32.76 ± 0.59<br>Late preterm (*n* = 13) : 35.31 ± 0.89 |
| Birth weight (g), *M* ± *SD*. | 3431.33 ± 506.55<br>Imputed (*n* = 1)<sup>a</sup> | 2243.24 ± 450.99 |
| (Corrected) Age at test (months), *M* ± *SD* | 6.13 ± 0.21 | 6.09 ± 0.19 |
| Sex: female / male | 12 / 18 | 6 / 11 |
| EPDS<sup>76</sup> score, *M* ± *SD* | 6.47 ± 3.37 | 5.59 ± 3.86 |
| IBQ<sup>77</sup> score, *M* ± *SD* |  |  |
| - Activity level | 4.09 ± 0.79 | 3.97 ± 1.00 |
| - Distress to limitations | 3.39 ± 0.66 | 3.33 ± 0.65 |
| - Distress and latency to intense stimulus | 2.45 ± 0.82 | 2.42 ± 0.64 |
| - Duration of orienting | 3.72 ± 1.09 | 4.00 ± 0.73 |
| - Smiling and laughter | 4.85 ± 0.94 | 4.69 ± 0.96 |
| - Soothability | 5.32 ± 1.05 | 5.10 ± 1.00 |
| PSI<sup>78</sup> score, *M* ± *SD* | 66.43 ± 11.55 | 66.82 ± 15.48 |
| SES<sup>79</sup> mother’s score, *M* ± *SD* | 1.33 ± 0.88 | 1.94 ± 1.39 |
| SES<sup>79</sup> father’s score, *M* ± *SD* | 1.90 ± 1.24<br>Imputed (*n* = 1)<sup>b</sup> | 1.59 ± 1.33<br>Imputed (*n* = 1)<sup>b</sup> |

The Cantonal Research Ethics Committee (CCER, BASEC No. 2022-02154) approved the protocol, and parents gave written informed consent before their infant took part, in accordance with the Declaration of Helsinki. Families received a small gift for their infant as a token of appreciation, and travel expenses were reimbursed on request; no other compensation was provided.

Among the 52 infants initially enrolled, 5 were excluded due to fussiness or technical issues during data collection (3 full-term and 2 preterm), yielding a final analytic sample of 47 infants (30 full-term and 17 preterm). For a subset of participants (*n* = 6), the experimental session could not be completed, resulting in partial datasets.

#### 1.2. Demographic variables

Demographic variables were compared between groups before the main analyses (cf. Table S1). The two missing paternal SES scores were replaced by the median of the infant’s group, and the single missing birth weight was predicted from gestational age by a regression fitted within the corresponding group, since the two variables are strongly associated within each group. Normality was examined in each group with the Shapiro-Wilk test, and continuous variables were compared with the Wilcoxon rank-sum test whenever normality was rejected in at least one group, and with Welch’s *t*-test otherwise. Sex distribution was compared with a chi-square test of independence, Fisher’s exact test being computed as a check. As expected, the groups differed in gestational age at birth and birth weight, two variables that define prematurity and that could therefore not be modeled independently of group. They did not differ in corrected age at testing, sex distribution, maternal mental well-being, infant temperament, or parental socioeconomic status, so that the group differences reported below are unlikely to reflect demographic, temperamental, or socioeconomic imbalances between the samples. Infant sex, recorded for every participant (cf. Table 1), was also screened as a model covariate and did not improve model fit (see Random effects and covariates).

### Method Details

#### 1.3. Overview of the design

Data were collected in the same project as a companion report^43^, which analyzed the term infants of the present sample to address a different question. The paradigm also contained trials with an omitted syllable, designed to probe temporal prediction and analyzed separately. Only sequences free of omissions are examined here.

#### 1.4. Voice recordings and stimuli

About a week before the visit, each mother sent back her recording, made after receiving written guidance and a sample file. She recorded herself at home, in a quiet room, with her own mobile phone held roughly 10 cm from her mouth, with neither the experimenters nor the infant present. Every syllable (/ba/, /ga/, /da/) was produced in three consecutive series of four repetitions, giving 12 tokens per syllable, and mothers were asked to follow the example and address their infant as they would in a calm moment to elicit infant-directed speech. The unfamiliar voice heard by a given infant came from the recording of the mother whose infant had been tested immediately before. All voices were thus recorded in the same way, and none was known to the infant who heard it. The very first participant heard, as an unfamiliar voice, a mother recorded for the pilot study under the same instructions.

Syllables were cut from the maternal recordings in Audacity v3.3–3.7^64^. Each syllable lasted 150 ms and began 20 ms before vowel onset, preserving the consonant. Syllables were exported as mono WAV files (48 kHz, 16-bit PCM), resampled to 44.1 kHz with librosa v0.10.2^65^, given 20 ms onset and offset ramps with the Praat Vocal Toolkit^66–2025^ in Praat v6.3.10^67^, peak-normalized, and scaled to 70 dB. After visual and auditory screening, the eight most homogeneous tokens of each syllable (positions 1–8 or 5–12) were kept.

#### 1.5. Testing procedure

Upon arrival, the EEG cap was fitted, and the infant and accompanying parent(s) were then taken into the testing room (i.e., a Faraday cage). Throughout data acquisition, the infant remained seated on the parent’s lap approximately 1 meter from a screen. Auditory material was played back at 65 dBA from two loudspeakers at the infant’s eye level and 1.5 m from the seating position; visual stimuli were rendered on a 1920 × 1080 px screen. Stimulus presentation was controlled with PsychoPy v2023.2.3^68^. To guarantee precise temporal alignment between events and the EEG trace, 10 ms trigger pulses marking stimulus onsets and offsets were routed to the amplifier through a parallel port.

Recording began once the infant had reached a quiet, awake state, with a 3-minute resting-state acquisition period, after which the experimental task was launched. Brief pauses were granted whenever the infant became unsettled, and feeding breaks were introduced when needed. Testing was discontinued if the infant did not return to a calm, wakeful state. Parents were instructed to refrain from any movement or interaction with the infant during stimulus blocks, to preserve signal quality and avoid behavioral confounds. Between blocks, however, soothing was permitted, as was the use of a pacifier.

#### 1.6. Session structure and stimulus timing

The session comprised 12 blocks, four for each rhythmic condition (delta-rate stimulation, theta-rate stimulation, and a non-rhythmic control), in a pseudo-randomized order. Each block started with 4–6 familiarization trials, followed by 8–10 test trials. Four test trials per block contained an omitted syllable, representing 25%–33% of a block’s trials, depending on its length (12–16 trials). Only sequences that neither contained an omission nor came right after one (non-violation sequences) entered the present analyses. Each trial was made of four successive four-syllable sequences. Within a trial, a single syllable (/ba/, /ga/, or /da/) was spoken by a single voice, that of the infant’s mother or of the unfamiliar mother, but with different recorded tokens, whose order changed between sequences according to a fixed set of permutations. Syllable and voice changed pseudo-randomly from one trial to the next within each block.

In delta-rate sequences, syllables started every 500 ms (2 Hz; measured asynchrony: 502 ± 10 ms), and in theta-rate sequences every 250 ms (4 Hz; measured asynchrony: 251 ± 10 ms), yielding sequences of 1.66 s and 0.90 s, respectively. For non-rhythmic sequences, new onset times were generated for each infant by randomly drawing the four onsets, at 1 ms resolution, within a 2-s window identical to that of delta-rate sequences. Successive onsets had to be at least 250 ms apart, and the fourth onset had to occur no later than 1800 ms after the start of the window. The resulting inter-onset intervals ranged from about 240 to 1090 ms, and sequence durations ranged from 0.97 to 1.94 s, with every syllable ending no later than 1.95 s after window onset. Sequences of a trial were separated by 1.39 s of silence at the delta rate, 1.14 s at the theta rate, and 1.1–2.5 s in the non-rhythmic condition, and consecutive trials by about 2 s.

Colored geometric shapes were displayed during the auditory stimulation, each visual onset coinciding with the onset of an auditory sequence. Auditory and visual stimuli were paired pseudo-randomly across blocks and infants. Changing syllables and visual displays from block to block aimed to maintain infants’ engagement throughout the session and to ensure that effects could be attributed to the temporal structure of the input rather than to particular features of the stimuli.

#### 1.7. EEG recording and questionnaires

Continuous EEG was acquired throughout the experimental protocol using a 128-channel Geodesic Sensor Net (GSN; Electrical Geodesics, Inc. [EGI]™). For most infants, the signal was relayed to an ANT Neuro eego™ amplifier, while a smaller number of recordings used an EGI amplifier; the same EGI net was employed in both setups to allow direct comparability across systems. The sampling rate was 1000 Hz, and impedances remained below 50 kΩ.

Several questionnaire-based measures were collected before the experimental session. Parental socioeconomic status (SES) was indexed using the Largo scale^79^, maternal mental well-being was assessed through the PSI^78^ and the EPDS^76^, and infant temperament was characterized using the IBQ^77^. Distributions of these measures are summarized in Table 1.

#### 1.8. EEG processing

Preprocessing was carried out in MNE-Python v1.9.0^62^ under Python v3.12.3, with six scripts run in sequence (see key resources table). Recordings were imported from BrainVision format, channels were renamed following the EGI E-notation, the GSN-HydroCel-128 montage was applied, and the auxiliary bipolar channels were removed. EEG triggers and the trial-by-trial behavioral log were aligned by an iterative matching procedure that removed unmatched rows until the two streams agreed. The resulting table coded, for each syllable, its identity, the voice (mother, stranger), the rhythmic condition (DEL, THE, NR), the position of the sequence relative to omissions (non-violation, before, during, or after a violation), and the position of the syllable within its sequence (1–4). One event marked the beginning of every four-syllable sequence. Sequence integrity was verified by checking onset intervals against the expected rate (500 ms at the delta rate, 250 ms at the theta rate) and confirming 16 syllables per trial; after this, recordings were split into session files.

Twenty-three peripheral electrodes, identical for all infants (E1, E8, E25, E32, E43, E48, E49, E56, E63, E68, E73, E81, E88, E94, E99, E107, E113, E119, E120, and E125–E128), were discarded, so that 105 channels remained. Channels judged noisy or flat on the raw traces and on the Welch power spectrum (0–150 Hz) were interpolated with spherical splines for each infant, and data were re-referenced to the common average. Signals were filtered between 1.5 and 100 Hz (zero-phase FIR filter, Hamming window), followed by a 50 Hz notch filter. Epochs from −2 to +7 s around sequence onset were extracted, epochs exceeding 5000 µV peak to peak being provisionally excluded, and downsampled to 250 Hz.

Independent component analysis was then run on these epochs with the extended Picard algorithm (python-picard v0.8^63^, 15 components, random_state = 20, max_iter = 10,000, decim = 4), and components reflecting ocular, cardiac, muscular, or sucking activity were removed after inspection of their topographies and time courses.

The cleaned epochs of all sessions of an infant were concatenated, and the behavioral information was realigned to the remaining epochs (epoch drop log and a combined timestamp-identity key), so that each epoch kept its condition labels. A copy of the epochs filtered between 1.5 and 30 Hz was produced for broadband analyses not reported here, and time-frequency representations were computed on the epochs that had not received this second filter.

#### 1.9. Time-frequency analysis

Multitaper convolution was used to obtain time-frequency representations of every channel and epoch between 2 and 98 Hz (2 Hz resolution), with n_cycles = f/2 and time_bandwidth = 2 for frequencies up to 40 Hz and n_cycles = f/5 and time_bandwidth = 6 for higher frequencies, and power was expressed in decibels (10 × log). After artifact rejection (see below), epochs were cropped to −1 to +2 s and baseline-corrected by subtracting, for each frequency, the mean power between −250 and −50 ms before sequence onset. The analysis targeted the delta (2–3 Hz) and theta (3–5 Hz) bands, both selected a priori based on previous infant EEG work. Mean power was extracted in the baseline window and in four consecutive post-onset windows (T1–T4), each as long as one syllable period of the condition: 0–500, 500–1000, 1000–1500, and 1500–2000 ms for delta-rate sequences, and 0–250, 250–500, 500–750, and 750–1000 ms for theta-rate sequences. Because syllables in non-rhythmic sequences did not follow a regular period, these sequences were split into the four 500-ms windows used for delta-rate sequences, which span the entire sequence.

#### 1.10. Artifact rejection

Epochs were rejected based on their peak-to-peak amplitude in two stages. The first, applied at epoching, excluded epochs above 5000 µV. The second relied on a threshold estimated from the data: peak-to-peak amplitudes of all infants, channels, and epochs (−500 to +2000 ms) were pooled, values outside the physiological range (below 1 µV or above 1000 µV) were discarded, and the threshold was set at the median ± 5 SD of the remaining distribution. This lenient criterion aimed to preserve as much data as possible from infant recordings. It was applied channel by channel, removing any epoch that exceeded the threshold, along with its time-frequency representation, only for the channel concerned; it was then applied again at the final stage of dataset construction. Epochs were then sorted by rhythmic condition, voice, and position relative to omissions, non-violation sequences being pooled with sequences that preceded an omission.

Depending on the number of blocks completed, the mean number of sequences presented per condition (rhythm × voice) ranged from 75.1 to 84.3 across conditions (individual values: 19–103). Overall, 7.9% of the presented sequences were rejected (term: 7.8%, preterm: 8.0%), mostly through the exclusion of whole epochs (7.0%), with the channel-wise criterion accounting for the remainder. Averaged over electrodes, the mean number of epochs retained per infant and condition ranged from 70.0 to 78.8 across conditions (individual values: 11.0– 96.9). Term and preterm infants did not differ in the total number of sequences presented (term: *M* = 496.7, *SD* = 71.4, preterm: *M* = 474.9, *SD* = 124.9, W = 263, *p* = .8675), in the number of epochs retained per condition (term: *M* = 76.3, *SD* = 13.3, preterm: *M* = 72.8, *SD* = 20.8, W = 275, *p* = .6659), or in the rejection rate (term: *M* = 8.0%, *SD* = 7.0, preterm: *M* = 8.3%, *SD* = 8.9, W = 259, *p* = .9382, Wilcoxon rank-sum tests).

#### 1.11. Region-of-interest definition

Following previous infant EEG studies^80–82^, eight a priori regions of interest (ROIs), each including 6 to 10 electrodes, were used: left frontal (LF: E12, E19, E20, E23, E24, E26, E27, E28, E33, E34), right frontal (RF: E2, E3, E4, E5, E116, E117, E118, E122, E123, E124), left temporal (LT: E35, E39, E40, E41, E45, E46, E50), right temporal (RT: E101, E102, E103, E108, E109, E110, E115), central (CEN: E7, E13, E31, E54, E55, E79, E80, E106, E112), left parietal (LP: E47, E51, E52, E53, E59, E60), right parietal (RP: E85, E86, E91, E92, E97, E98), and posterior (POS: E61, E62, E67, E72, E77, E78). Electrode E26 was absent from the final dataset, so that LF comprised nine electrodes in the analyses. Difference time-frequency maps between conditions were computed for each ROI in the two bands of interest for visual inspection.

#### 1.12. Dataset construction

Single-infant files were pooled across the sample, separately for each voice × rhythm combination. A final table then stored one row per surviving epoch, channel, band, and time window, with the mean power in that window and the condition labels (rhythmic condition, voice, frequency band, channel, position relative to omissions, and block order). All models used these single-epoch values. Epoch averages served only for the figures, over −250 ms to +2 s (delta-rate and non-rhythmic sequences) or −250 ms to +1 s (theta-rate sequences).

### Quantification and Statistical Analysis

#### 1.13. Statistical models

Analyses were run in R v4.5.1^69^ with glmmTMB v1.1.13^70^. The dependent variable was baseline-corrected power (dB) in the delta and theta bands, modeled with Gaussian generalized linear mixed models (GLMMs; identity link). Each model was fitted once on the eight ROIs, with ROI entered as a fixed factor crossed with all other fixed effects (10,159,904 single-epoch observations). Frequency band (FRQ: delta, theta) was a fixed factor, the four post-onset windows (T1–T4) were pooled, and only non-violation sequences were included. The fixed-effect design matrix was stored in sparse form to keep memory requirements manageable. Models were first fitted with the nlminb optimizer. When it ended with a false convergence, the fit was restarted from the parameters reached and, if needed, refit with the BFGS algorithm, a refit being kept only if it converged with a positive definite Hessian and an AIC no higher than that of the initial fit, which was the case for prediction (b).

Two predictions were evaluated:

a. neural tracking would be jointly shaped by rhythmic context (RHY: DEL, THE, or NR stimulations), frequency band (FRQ: Delta and Theta), and prematurity status (GROUP: term vs. preterm), and was operationalized as: Power ∼ RHY × FRQ × GROUP × BLK_ORD × ROI + (1|PAT) + (1|CH).
b. this prediction was extended by introducing voice familiarity (VOICE: Mother vs. Stranger) as an additional fixed factor, yielding: Power ∼ RHY × FRQ × VOICE × GROUP × BLK_ORD × ROI + (1|PAT) + (1|CH).

Both predictions were formulated on theoretical grounds before any model was fitted, although the study was not preregistered. Tests of these predictions are treated as confirmatory and all other analyses as exploratory.

Because the question concerns how tracking evolves during exposure, both predictions were expressed as slopes, with block order (BLK_ORD) entering as a continuous predictor in interaction with all other fixed effects. To make slopes specific to each rhythmic condition, blocks were renumbered from 1 to 4 within each infant and rhythmic condition, so that BLK_ORD indexes the successive blocks of a given rhythm.

#### 1.14. Random effects and covariates

The random-effects structure was selected by comparing, for each prediction, four models with the same fixed effects (RHY × FRQ × GROUP × ROI, with VOICE added for prediction b): no random intercept, a participant intercept (1|PAT), an electrode intercept (1|CH), or both. Convergence of each candidate was checked (optimizer code, positive-definite Hessian, finite standard errors, and non-degenerate variances), with the refitting procedure described above. Including both intercepts gave the lowest AIC for both predictions; removing the participant intercept increased AIC by more than 5,600, and removing the electrode intercept by 58.7. The structure (1|PAT) + (1|CH) was therefore used in all models.

Twelve candidate covariates were then screened: infant sex (SEX), maternal depressive symptoms (EPDS), parenting stress (PSI), six IBQ subscales (IBQ_AL, IBQ_DL, IBQ_DI, IBQ_DO, IBQ_SL, IBQ_S), maternal and paternal socioeconomic status (SES_M, SES_P, missing values replaced by the group median), and block order (BLK_ORD). For each prediction, the base model was compared with twelve models, each adding one covariate, using AIC and likelihood-ratio tests. A covariate was retained only if the likelihood-ratio test was significant (*p* < .05) and the AIC comparison agreed. Only BLK_ORD, coded from 1 to 4 within each rhythmic condition, met this criterion (prediction a: χ²(1) = 19.07, *p* < .0001, ΔAIC = 17.1, prediction b: χ²(1) = 16.89, *p* < .0001, ΔAIC = 14.9), and none of the eleven other candidates improved the fit (all *ps* ≥ .2271). BLK_ORD was therefore the only covariate in the final models. Together with the absence of group differences in background variables, this suggests that the GROUP effects reported here relate to prematurity rather than to demographic, psychosocial, temperamental, or socioeconomic differences between the samples.

#### 1.15. Omnibus test and confirmatory contrasts

Each fixed effect and interaction was evaluated with a Type II Wald χ² test (car v3.1.3^71^). These omnibus tests were not corrected for multiple comparisons, as each term was tested a single time in one model covering all ROIs. The rule for follow-up tests was fixed before they were computed: if an interaction including ROI was significant, slopes and contrasts were estimated within each ROI, and if only the corresponding interaction without ROI was significant, they were averaged over ROIs. Since the interactions including ROI were significant (Table S2), all follow-up estimates are given per ROI. Marginal slopes of BLK_ORD were obtained with emmeans v2.0.0^72^ (emtrends) for each group, rhythmic condition, band, and ROI (prediction a), and additionally for each voice (prediction b). Each slope therefore indicates whether tracking increased, decreased, or remained stable over successive blocks of a given rhythm. Term and preterm slopes were compared by pairwise contrasts (Term − Preterm).

The distinction between canonical tuning (stimulation rate matching the band, e.g., delta-rate stimulation in the delta band) and cross-frequency tuning (mismatch, e.g., delta-rate stimulation in the theta band) was part of the rationale from the outset. The three rhythmic conditions were nevertheless entered together in the confirmatory models, the non-rhythmic condition serving as the reference against which rhythm-specific effects are defined. A slope change specific to one rhythmic condition is therefore unlikely to reflect a uniform drift over the session, and contrasts for the non-rhythmic condition are reported alongside the canonical and cross-frequency contrasts.

*p* values and confidence intervals of slope contrasts were adjusted with the multivariate t (mvt) method, which provides simultaneous 95% confidence intervals, with one family per frequency band including all ROIs (24 Term − Preterm contrasts per band for prediction a and 48 for prediction b). The random seed of the mvt computation was fixed. Slopes of each group are reported with unadjusted 95% confidence intervals and without a test against zero. All tests were two-sided, with α = .05.

##### 1.15.1. Exploratory analyses

Three exploratory analyses followed. First, block-wise differences (Δ BLK_ORD, block 4 minus block 1 and block 4 minus block 2) were computed for the tuning and cross-frequency contrasts that were significant in the confirmatory step (cf. Tables S3 and S4). Because BLK_ORD is continuous, these quantities are three and two times the slope difference, respectively, and share its *z* and *p* values. They are reported only to express the effects over the whole session.

Second, frequency selectivity was indexed by comparing, within each response band, the slope for the matching rhythm with the slope for the non-matching rhythm: delta-rate minus theta-rate slope in the delta band, and theta-rate minus delta-rate slope in the theta band.

Because it contrasts two slopes, this index describes how the gap between matched and mismatched responses evolved during the session, with positive values indicating that it widened and negative values indicating that it narrowed. The non-rhythmic condition was not part of the index. For each ROI and band, selectivity in term infants, selectivity in preterm infants, and their difference were computed as linear combinations of the model slopes, with standard errors accounting for the covariance between slopes. *p*-values and confidence intervals were mvt-adjusted with one family per band (three quantities × eight ROIs, 24 contrasts; Table S5).

Third, to examine more directly how voice shaped the group difference in canonical delta tuning, a reduced GLMM was fitted to the delta stimulation/Delta observations of all eight ROIs (1,678,516 observations): Power ∼ VOICE × GROUP × BLK_ORD × ROI + (1|PAT) + (1|CH). In this model, the VOICE × GROUP × BLK_ORD interaction is a single term that can be tested directly. Two sets of contrasts were derived from it, each mvt-adjusted as a single family across ROIs: Term − Preterm slope differences for each voice (16 contrasts), and the difference between voices in the Term − Preterm gap at each block (32 contrasts). Results are reported in Figure S1 and Tables S10–S14.

#### 1.16. Effect sizes and model diagnostics

Model fit is described by the marginal and conditional R² (R²m, R²c)^83^, the log-likelihood, and the convergence status (Tables S6, S7, and S13). R² values were computed from the variance components of each model, because the performance package v0.15.3^73^ does not handle sparse design matrices; the two computations gave identical values on simulated data. As expected for single-epoch spectral power, the models explained little variance (Prediction a: R²m = 0.0003, R²c = 0.0009; Prediction b: R²m = 0.0005, R²c = 0.0011; reduced model: R²m = 0.0004, R²c = 0.0019). Slopes and contrasts are reported as raw model estimates, in dB per block, with asymptotic standard errors and 95% Wald confidence intervals (unadjusted for group slopes, simultaneous for contrasts) and without further standardization (Tables S3, S4, S8, S9, S11, S12, and S14). Residuals of each model were examined with DHARMa v0.4.7^74^ (250 simulations) for uniformity, dispersion, and outliers. Because a simulation matrix of about 10 million observations would exceed available memory, each model was simulated on all observations, and scaled residuals were computed on a random subsample of 200,000 observations. Dispersion was adequate in all three models (dispersion ratio = 1.000, 1.003, and 0.996 for Prediction a, Prediction b, and the reduced model, all *p*s ≥ .2960). With 200,000 residuals, the Kolmogorov-Smirnov and outlier tests detect even negligible departures (all *p*s < .0001), so the size of the deviations is the relevant information: the distance between the observed and expected residual distributions was small (*D* = 0.016, 0.014, and 0.019), outliers were only slightly more frequent than expected (1.01% to 1.04% vs. 0.80%), and residual plots showed no systematic pattern apart from a slight upward deviation of the median residual at the highest predicted values. Figures were produced with ggplot2 v4.0.0^75^. No statistical method was used to determine sample size in advance.

## Resource Availability

### Lead contact

Requests for further information and resources should be directed to and will be fulfilled by the lead contact, Isabelle Rambosson.

### Materials availability

This study did not generate new materials. The auditory stimuli are recordings of participants’ mothers and are subject to the same consent restrictions as the EEG data.

### Data and code availability

The EEG data cannot be deposited in a public repository because of restrictions stated in the informed consent approved by the Cantonal Research Ethics Committee (BASEC No. 2022-02154). De-identified data will be shared by the lead contact with qualified researchers upon reasonable request, subject to ethics committee approval.

All original code (Python preprocessing pipeline and R analysis scripts) has been deposited at Yareta and is publicly available at https://doi.org/10.26037/yareta:4dtcmgkw4jdwpmawpzcdt4byrq.

Any additional information required to reanalyze the data reported in this paper is available from the lead contact upon request.

## Author contributions

I.R., D.B., C.K., D.G. and M.F. conceived the study. I.R., D.B., L.C., D.G. and M.F. developed the methodology. I.R., D.B., F.B.-M. and M.F. carried out the investigation. I.R. performed the formal analysis and prepared the visualizations. L.C., D.G. and M.F. validated the results. D.B., L.C., D.G. and M.F. supervised the work. D.G. and M.F. acquired funding.

I.R. and M.F. wrote the original draft. I.R., D.B., F.B.-M., L.C., D.G. and M.F. reviewed and edited the manuscript. D.G. and M.F. jointly supervised this work. All authors approved the final version of the manuscript.

## Acknowledgement

We thank the Arcade Sages-Femmes Association (Geneva) and Dr. Marie Janaillac (specialist in pediatrics and neonatology, Clinique des Grangettes) for helping recruit participants. We also thank Emmanuelle Maillard, Léa Ghezzi, and Nina Luna Marbehant (students, University of Geneva) for their contribution to data collection, as well as Maïté Fontela and Lilou Dehondt (students, University of Geneva) for their involvement in data collection, preprocessing, and processing.

